# Bayesian inference of lineage trees by joint analysis of single-cell multimodal lineage-tracing data with BiLinT

**DOI:** 10.1101/2025.09.21.677669

**Authors:** Ziwei Chen, Bingwei Zhang, Linrui Tang, Fuzhou Gong, Lin Wan, Liang Ma

## Abstract

The advent of single-cell lineage-tracing technologies has enabled simultaneous measurement of gene expression and lineage barcodes. However, integrating these modalities for high-resolution lineage reconstruction has remained challenging due to limitations of methods that analyze modalities separately. In response, we present BiLinT, a Bayesian framework for reconstructing high-resolution cell lineage trees by jointly modeling single-cell multimodal lineage-tracing data. BiLinT integrates lineage barcode evolution (modeled via a continuous-time Markov chain) and gene expression dynamics (modeled via an Ornstein-Uhlenbeck process) within a unified probabilistic model. Applications to synthetic and real datasets demonstrate improved accuracy and reveal developmental fate biases.

## Introduction

Understanding lineage dynamics and the associated cellular state-transition trajectories during the formation of tissues, organs, and whole organisms is fundamental to developmental biology (Kalhor et al. 2018; Wagner et al. 2018; Chan et al. 2019; C Chen et al. 2022) and cancer progression (Navin and Hicks 2010; Caiado et al. 2019; Yang et al. 2022; Jones, Yang, et al. 2023; Lamprecht et al. 2017; Pai et al. 2023). Due to limitations in directly observing cell division processes within intact tissues or whole organisms, recent advances in dynamic lineage tracing have introduced the use of genome editing technologies, particularly CRISPR/Cas9, to induce heritable mutations at predefined genomic target sites (Hsu et al. 2014; Doudna and Charpentier 2014; Jiang and Doudna 2017). This approach enables the manipulation of the number of target sites and genomic editing parameters to obtain richer phylogenetic signals. When combined with single-cell sequencing, CRISPR-based lineage tracing enables the simultaneous capture of clonal relationships and transcriptomic profiles for thousands of individual cells (McKenna and Gagnon 2019; Sankaran et al. 2022; Raj 2023; Chehelgerdi et al. 2024). Techniques such as scGESTALT (Raj et al. 2018), LINNAEUS (Spanjaard et al. 2018), iTracer (Z He et al. 2022), CREST (Xie et al. 2023), and CARLIN/DARLIN (Bowling et al. 2020; L Li et al. 2023) provide effective pipelines for tracking cellular lineages and the associated changes in cellular states throughout cellular development or disease progression.

Despite these technological advances, reconstructing accurate and high-resolution cellular developmental trajectories from single-cell lineage-tracing data remains a significant challenge. The quality and characteristics of CRISPR/Cas9-based barcode data impose fundamental limitations on lineage reconstruction accuracy (Salvador-Martínez et al. 2019). Factors such as a limited number of target sites (Pan, H Li, Putta, et al. 2023), frequent dropout events (Pan, H Li, Putta, et al. 2023; Mai et al. 2024; Seidel and Stadler 2022), and the non-uniform distribution of mutations across loci (Pan, H Li, and X Zhang 2022; Salvador-Martínez et al. 2021; Pan, H Li, Putta, et al. 2023) hinder the generation of high-resolution lineage trees.

A number of computational methods, such as Cassiopeia (Jones, Khodaverdian, et al. 2020), DCLEAR (Gong, Kim, et al. 2022), Startle (Sashittal et al. 2023), and TiDeTree (Seidel and Stadler 2022), have been developed to infer lineage trees solely from barcode data. These approaches can be broadly categorized into four types: distance-based methods (Gong, Kim, et al. 2022; R Wang et al. 2023), maximum parsimony methods (Jones, Khodaverdian, et al. 2020; Sashittal et al. 2023), maximum likelihood methods (Zafar et al. 2020; Feng et al. 2021; Mai et al. 2024), and Bayesian inference methods (Seidel and Stadler 2022). While maximum likelihood and Bayesian methods typically offer higher accuracy, their exclusive reliance on barcode data imposes fundamental limitations (Hasegawa and M Fujiwara 1993; Gadagkar and Kumar 2005; Swofford et al. 2001). In particular, the ambiguity caused by barcode dropouts and limited target diversity can result in multiple cells sharing identical barcode states, leading to shallow trees with multifurcated internal nodes that obscure developmental resolution (Pan, H Li, and X Zhang 2022; Salvador-Martínez et al. 2021; Gong, Granados, et al. 2021).

To enhance lineage resolution beyond barcode data alone, recent methods such as LinTIMaT (Zafar et al. 2020) and LinRace (Pan, H Li, Putta, et al. 2023) incorporate gene expression information. LinTIMaT introduces a combined likelihood framework under the assumption that transcriptionally similar cells are likely to share similar barcode states and thus reside near each other in the lineage tree. However, this assumption does not always hold in practice: cells with nearly identical transcriptomes may appear on distant branches, and conversely, cells within the same subtree can exhibit substantial transcriptional heterogeneity (Chan et al. 2019; Raj et al. 2018; Pan, H Li, Putta, et al. 2023). Consistent with this, LinTIMaT does not always outperform barcode-only approaches in synthetic benchmarks (Pan, H Li, and X Zhang 2022). LinRace seeks to address this challenge by introducing an asymmetric cell division model and adopting a two-step strategy in which a fixed barcode-based backbone tree is first inferred and then refined using transcriptomic similarity. While intuitive, this sequential design inherits two important limitations (Bastide et al. 2021). First, it treats the backbone tree as known and fixed, thereby failing to propagate the considerable uncertainty inherent in reconstructing lineage relationships from barcode data. Second, by conditioning refinement strictly on the initial backbone, the method does not fully exploit gene expression information during tree reconstruction, but only afterward in subtree adjustment. These restrictions can limit its ability to capture global lineage structure and fully integrate the complementary strengths of barcode and transcriptomic signals.

To address these limitations, we introduce BiLinT, a Bi-process-based Bayesian Lineage Tree reconstruction method that simultaneously models lineage barcodes with a continuous-time Markov chain and gene expression dynamics with an Ornstein–Uhlenbeck (OU) process (Fig. 1). Compared to local asymmetric models, the OU process provides a global, continuous representation of gene expression dynamics along lineages, capturing both stochastic fluctuations and stabilizing selection. This joint modeling enables BiLinT to reconstruct either a complete single-cell phylogeny or clone-level trees, grouping cells by shared barcode states and latent transcriptional identities to capture both shared ancestry and convergent phenotypes.

**Figure 1.**
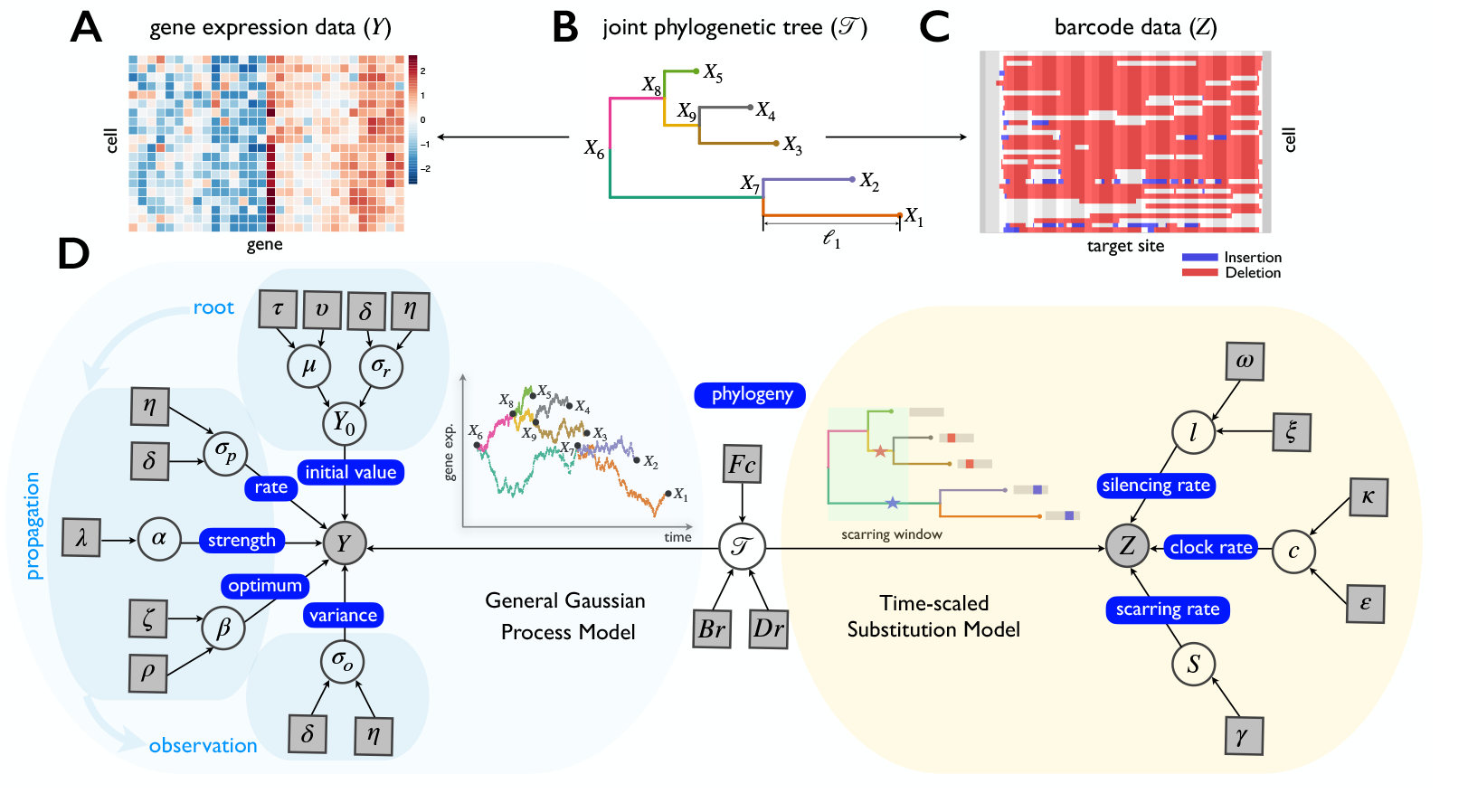
Overview of the computational framework of BiLinT that integrates scRNA-seq data and lineage barcode data for the lineage-tracing process. (A) scRNA-seq gene expression data (***Y***), where each row indicates a cell and each column represents a gene. (B) The latent joint phylogenetic tree (𝒯). (C) Lineage barcode data (***Z***), where each row represents one cell and each column is a target site. (D) The probabilistic graphical model shows the dependencies among parameters, where the shaded nodes stand for observed or fixed values, the unshaded nodes represent the latent parameters.

The framework explicitly accounts for barcode complexities, such as silent states, variable editing efficiencies, and parallel mutations, as well as measurement noise inherent in gene expression data. Efficient inference via Markov Chain Monte Carlo (MCMC) enables scalable parameter estimation and tree reconstruction with linear complexity. Benchmarking on synthetic datasets confirms BiLinT’s superior accuracy in cell- and clone-level tree reconstruction. Applied to developing mouse ventral midbrain (vMB) and hematopoietic stem cells, BiLinT uncovers spatially distinct lineages and fate biases, demonstrating its capability to resolve detailed lineage structures from complex multimodal datasets.

## Results

### Overview of BiLinT

We present a unified Bayesian framework integrating single-cell RNA sequencing (scRNA-seq) (Fig. 1A) and single-cell lineage barcode data (Fig. 1C) for joint phylogenetic analysis of developmental processes using snapshot profiles of both cell states and cell lineages.

We assume a hidden phylogenetic tree 𝒯 that captures the ancestral relationships among *N* observed cells, each mapped to the tips of the tree (Fig. 1B). Given this tree structure, BiLinT jointly analyzes barcode data *Z* and gene expression data *Y*, assuming that the two data types evolve independently along the branches conditioned on the phylogenetic tree (Fig. 1D).

BiLinT models the barcode data ***Z*** using a time-dependent substitution model adapted to CRISPR-Cas9 editing events. Based on the foundation of TiDeTree (Seidel and Stadler 2022), this substitution model reflects the time-scaled nature of lineage trees, where branch lengths correspond to absolute durations of cell division. It also accounts for parallel mutations, which are independent mutations at the same site across different branches, thereby broadening the space of plausible tree topologies. Additionally, the model addresses silent loci and variability in editing efficiency, ensuring a close match to real barcode data.

Simultaneously, BiLinT models the gene expression data ***Y*** by a continuous-time, continuous-state Markov process that evolves along the tree branches. Specifically, we adopt the OU process to model gene expression dynamics (Dimayacyac et al. 2023; Rohlfs et al. 2014; Bedford and Hartl 2009; J Chen et al. 2019; Rattray et al. 2019; Hirsch et al. 2025; Bastide et al. 2021). As a generalization of Brownian motion, the OU process has bounded stationary variance, making it suitable for modeling gene expression dynamics that stabilize over time. To account for measurement noise in scRNA-seq data, an observation model links the latent state at the tip of the embedded process with the observed expression values.

Let *ϕ* represent all parameters related solely to the barcode evolution model and *θ* denote the parameters associated with the continuous process of gene expression. BiLinT assumes conditional independence of the two processes given the tree and aims to infer the posterior distribution of the tree structure and parameters, expressed as:

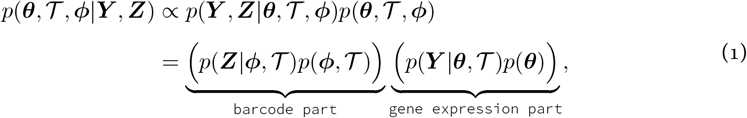

where *p*(***Y***, ***Z***|*θ*, 𝒯, **φ**) is the likelihood of observing the data given the parameters and tree, and *p*(*θ*, 𝒯, ***φ***) is the prior over the parameters and tree structure. By assuming conditional independence between the barcode and gene expression data given the tree 𝒯, the likelihood decomposes into two components: *p*(***Z***| ***φ***, 𝒯), representing the likelihood of barcode data, and *p*(***Y*** |*θ*, 𝒯), representing the likelihood of gene expression data. This decomposition simplifies the joint inference while retaining the contributions from both data types. To infer the posterior distribution over the tree and parameters, BiLinT employs an MCMC algorithm.

### BiLinT reconstructs joint trees with high accuracy and efficiency on simulation datasets

We benchmark BiLinT against state-of-the-art methods for tree inference based on lineage-tracing data—including LinRace, LinTIMaT, Cassiopeia, DCLEAR, and the Neighbor-Joining (NJ) method—across three groups of simulation datasets, totaling 121 runs under diverse conditions.

To evaluate the impact of different gene expression models, we simulate expression data using two approaches: the asymmetric cell division model implemented in TedSim (Pan, H Li, and X Zhang 2022) (Simulation 1; 90 datasets) and our General Gaussian Process Model (Simulation 2; 30 datasets). Barcode data are generated using our time-scaled substitution model in both cases. Since TedSim is restricted to balanced tree topologies, Simulation 1 uses balanced phylogenetic trees, whereas Simulation 2 employs trees drawn from a birth–death–sampling process, allowing for randomly shaped, unbalanced trees. To further highlight the benefit of joint modeling, we design Simulation 3, a challenging scenario where barcode information is limited due to a small number of editable sites and frequent parallel mutations. This dataset includes eight clones arranged along an unbalanced tree, with gene expression simulated by the General Gaussian Process Model (Fig. 3A) and fixed barcode states listed in Table S1.

In Simulation 1, we systematically vary three key parameters—clock rate *c*, silencing rate *l*, and number of cells *N* —while holding others fixed. Across these conditions, BiLinT performs comparably to LinRace and outperforms LinTIMaT, Cassiopeia, DCLEAR, and NJ in terms of RF and SPR distances, which assess the topological and structural congruence between the reconstructed trees and the ground truth (Fig. S1-S3). All methods improve with higher clock rates (Fig. S1), but accuracy declines with increased silencing (Fig. S2) or larger cell numbers (Fig. S3). Notably, BiLinT consistently achieves superior accuracy in path-distance metrics due to its time-scaled framework.

In Simulation 2, we test BiLinT under more realistic, unbalanced tree structures, varying the number of cells *N*. BiLinT demonstrates robust performance across all metrics (Fig. 2 and S4) and consistently outperforms LinRace and other baseline methods, particularly in reconstructing cell trees.

**Figure 2.**
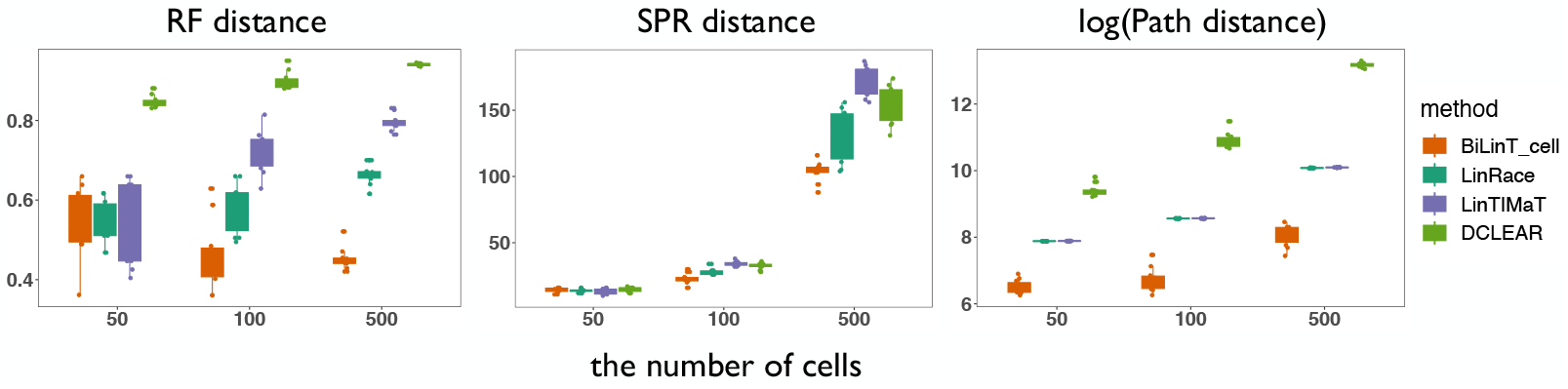
Comparison of performance among BiLinT, LinRace, LinTIMaT, and DCLEAR in reconstructing cellular trees using simulations generated by the BiLinT model with varying cell counts (Simulation 2). Since SPR distance is not applicable to clonal trees, results for the BiLinT clone tree and Cassiopeia clone/binary tree are not presented here. For comprehensive results including performance of BiLinT clone tree and Cassiopeia trees, please refer to Fig. S4.

**Figure 3.**
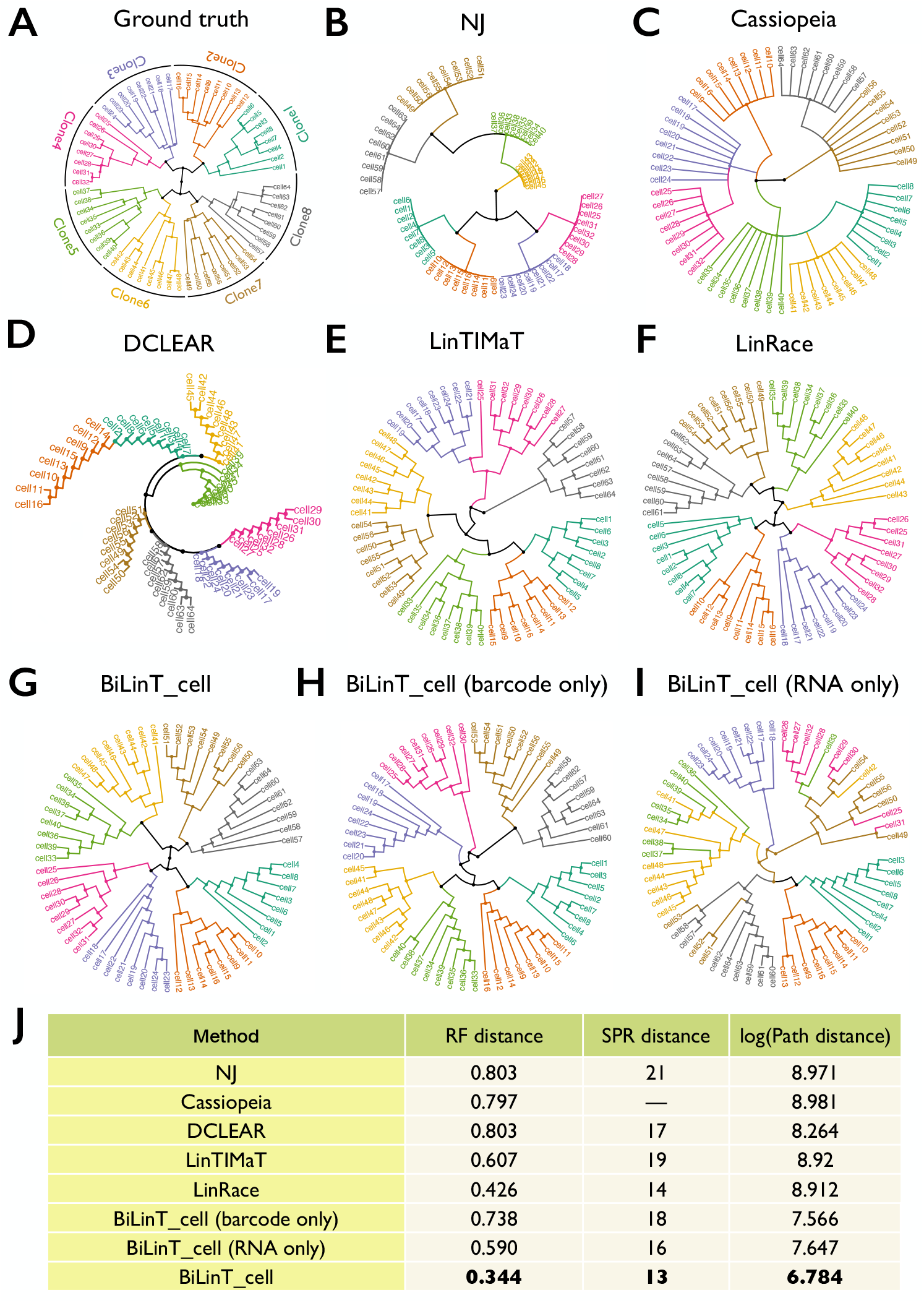
Comparison of reconstructed cellular trees and evaluation metrics under simulated conditions with limited target sites and parallel mutations (Simulation 3). (A) Ground truth tree. (B–D) Trees reconstructed using barcode-only methods: NJ, Cassiopeia, and DCLEAR, respectively. (E–G) Trees reconstructed using joint analysis of barcode and RNA data with LinTIMaT, LinRace, and BiLinT, respectively. (H–I) Trees reconstructed by BiLinT using barcode-only data and RNA-only data, respectively.

In Simulation 3, we compare methods under conditions where barcode data alone provide weak phylogenetic signal. We first use the NJ method, Cassiopeia, and DCLEAR to generate phylogenetic trees based solely on lineage barcode data (Fig. 3B–D). The NJ method organizes clones 5–8 in a hierarchical structure. In the Cassiopeia tree, clones 1, 5, and 6 are grouped on the same main branch, while clones 2–4 form a separate lineage. DCLEAR, on the other hand, reconstructs an unbalanced tree. Subsequently, we apply BiLinT, LinRace, and LinTIMaT to jointly infer cell trees by integrating barcode and scRNA-seq data (Fig. 3E–G). LinTIMaT places clone5 alongside clone1 and 2 in one main lineage, and incorrectly merges clone7 with clone6. LinRace uses the NJ tree as a fixed backbone for barcode data without accounting for uncertainties, which introduces biases that propagate into its final structure. In contrast, BiLinT successfully recovers the correct topological structure among all clones. Furthermore, BiLinT demonstrates superior performance across all three evaluation metrics, as shown in Fig. 3J. To further highlight the importance of joint modeling, we also test BiLinT using only barcode or only gene expression data (Fig. 3H–I). In both cases, BiLinT fails to reconstruct the correct clone relationships, resulting in significantly higher distances to the ground truth compared to the joint-inference model (Fig. 3J). These results underscore the value of BiLinT’s integrated approach in leveraging both data modalities for accurate lineage reconstruction.

### BiLinT reveals fate bias within developing HSCs

We next investigate lineage fate bias in hematopoietic development by applying BiLinT to a single-cell lineage-tracing dataset generated using the inducible Cas9 barcoding system DARLIN (L Li et al. 2023). In this system, doxycycline (Dox) is administered at embryonic day 17.0 (E17.0) to activate barcode editing, and Lin^−^ Kit^+^ cells are isolated from skull bone marrow nine weeks later. The DARLIN model uses terminal deoxynucleotidyl transferase (TdT) and 30 CRISPR target sites distributed across three barcode arrays (CA, TA, and RA), which exhibit near-complete editing (~100%) within 24 hours of Dox induction. Thus, the effective scarring window spans 1 day, reflecting the rapid editing kinetics of the barcode arrays, while the total developmental time extends to 64 days (from E17.0 to sample collection).

After filtering low-quality cells and clones using the CARLIN pipeline (Bowling et al. 2020) and retaining only robust clones with *≥* 15 cells, we obtained a dataset of 1,003 cells containing both lineage barcodes and transcriptomes. Cell types were annotated as hematopoietic stem cells (HSC), lymphoid-biased multipotent progenitors (LMPP), megakaryocyte progenitors (MkP), erythrocytes (Ery), neutrophils (Neu), monocytes (Mon), basophils (Baso), and dendritic cells (DC) (Fig. 4A). Using default parameters, BiLinT jointly inferred the phylogenetic tree by integrating barcode mutations and gene expression dynamics—modeled via a continuous-time substitution process and an OU process, respectively.

**Figure 4.**
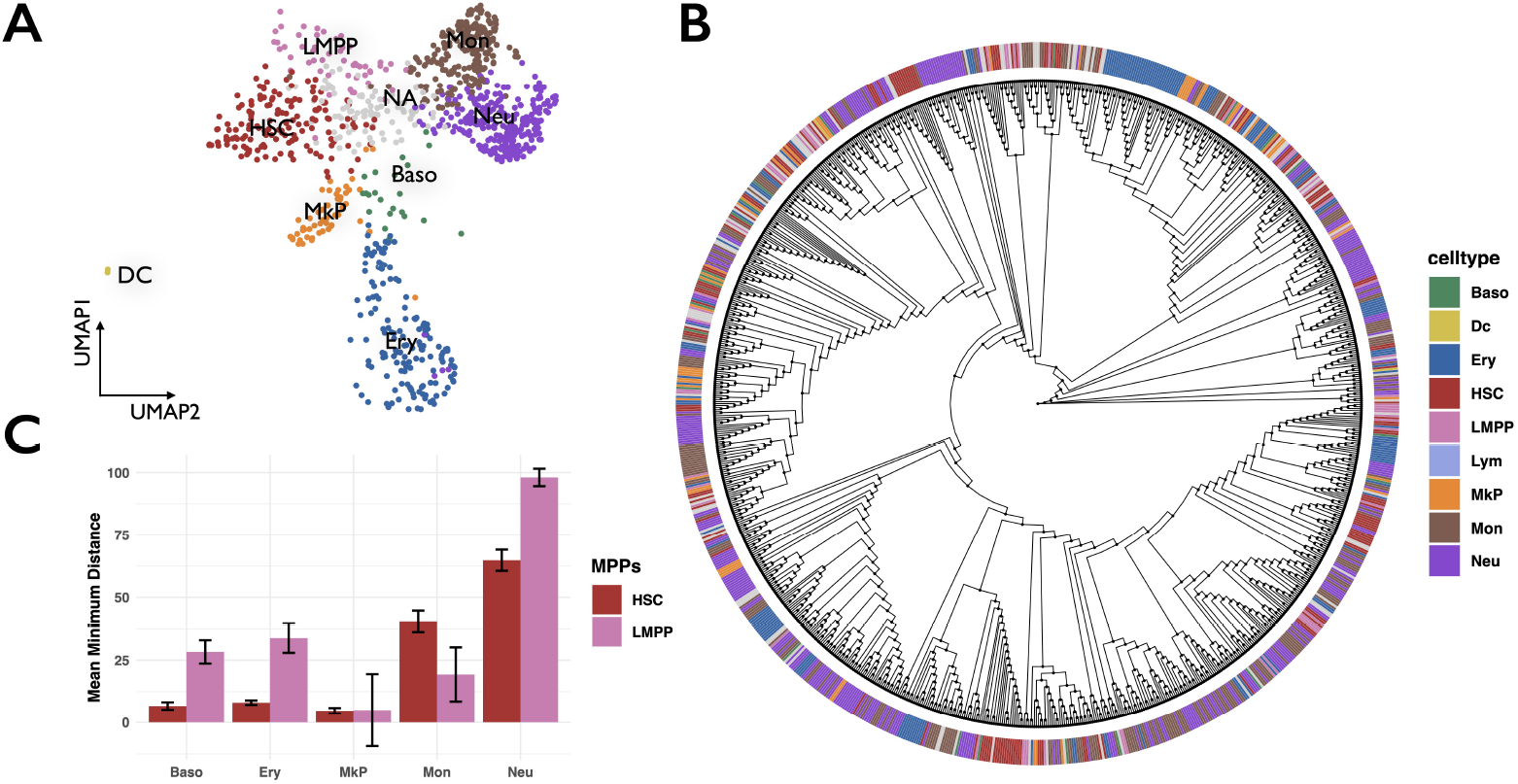
BiLinT reveals fate bias within developing HSCs on DARLIN datasets. (A) UMAP visualization of DARLIN scRNA-seq dataset. (B) BiLinT reconstructs cell lineage tree for the DARLIN dataset. (C) The mean of the minimum distances along BiLinT tree from HSCs and LMPPs to various differentiated cell types is calculated. Due to the limited number of DCs (only 2), this cell type is excluded from the analysis.

BiLinT reconstructs a unified lineage tree that reveals both clonal relationships and continuous differentiation trajectories (Fig. 4B). To assess fate bias, we compute the mean minimum distances from HSCs and LMPPs to each mature cell type along the BiLinT-inferred tree. The results show that HSCs are significantly closer to MkPs than to other lineages (*p*_*value*_ *<* 2.2 × 10^−16^, Wilcoxon rank-sum test), consistent with previous reports of direct HSC-to-MkP differentiation (AE Rodriguez-Fraticelli, Wolock, et al. 2018; AE Rodriguez-Fraticelli, Weinreb, et al. 2020; Nishikii et al. 2015; Roch et al. 2015; Carrelha et al. 2018; Morcos et al. 2022) (Fig. 4C). Similarly, LMPPs exhibit shorter distances to monocytes than to neutrophils (*p*_*value*_ *<* 2.2 × 10^−16^, Wilcoxon rank-sum test), suggesting a bias toward monocytic fate. These findings demonstrate BiLinT’s ability to resolve both discrete clonal ancestries and continuous differentiation paths. In particular, the OU process enables BiLinT to capture smooth transitions in gene expression along lineages, facilitating high-resolution inference of fate bias in complex developmental systems.

### BiLinT reconstructs the lineage landscape for the developing mouse ventral midbrain

We apply BiLinT to paired lineage barcode and transcriptomic data from En1-Cre-CREST mouse embryos at E11.5 and E15.5 to investigate neurodevelopment of ventral midbrain (vMB). In this system, Cre recombinase drives lineage barcoding between E8.0 and E8.5, and approximately 99% of barcode editing is completed by E8.5 (Xie et al. 2023). scRNA-seq data from barcoded cells are then collected at E11.5 and E15.5, capturing key neurogenic transitions. Accordingly, the effective barcoding window spans 0.5 days (E8.0–E8.5), with total developmental time extending to 3.5 days (E8.0–E11.5) and 7.5 days (E8.0–E15.5).

We preprocess the data and retain robust clones containing at least 15 cells at both time points. To compensate for the relatively low number of cells and clones at E11.5, we include three biological replicates. After filtering out low-quality cells and clones, the three E11.5 replicates contribute 52, 53, and 179 cells, distributed across 3, 3, and 8 clones, respectively. The single E15.5 replicate yields 1,561 cells within 48 clones. We apply BiLinT to jointly infer lineage trees by integrating both barcode information (V1 and V2 CREST barcodes) and gene expression data. Cell type diversity at each stage is shown in Fig. S5A,D. Hierarchical clustering of cell-type profiles in (Xie et al. 2023) reveals distinct spatially organized lineage groups, or principal lineages, including floor plate (FP), basal medial (BM), and basal lateral (BL) at E11.5, and additional alar lateral (AL) and mid-hindbrain boundary (MHB) domains emerging at E15.5.

BiLinT reconstructs comprehensive lineage trees by integrating barcode and transcriptomic information, enabling fine-grained resolution of developmental hierarchies. Within each reconstructed subtree or clone, BiLinT identifies both clonal expansion and transcriptional progression, with cells of the same type clustering closely and clear transitions from early progenitors and Rgl cells to neuroblasts and terminal neuron types (Fig. 5A–B, Fig. S5B–C). These transitions are not discrete but continuous, as captured by the OU process, which models gene expression dynamics as cells evolve along the tree. The OU framework allows BiLinT to infer smooth expression trajectories within each subtree, distinguishing between closely related but transcriptionally distinct cell states. Furthermore, BiLinT reveals that cells of the same type often appear across multiple subtrees, reflecting the “partial consistency” (Pan, H Li, Putta, et al. 2023) between transcriptomic similarity and barcode-derived clonal structure—an inherent characteristic of single-cell lineage data.

**Figure 5.**
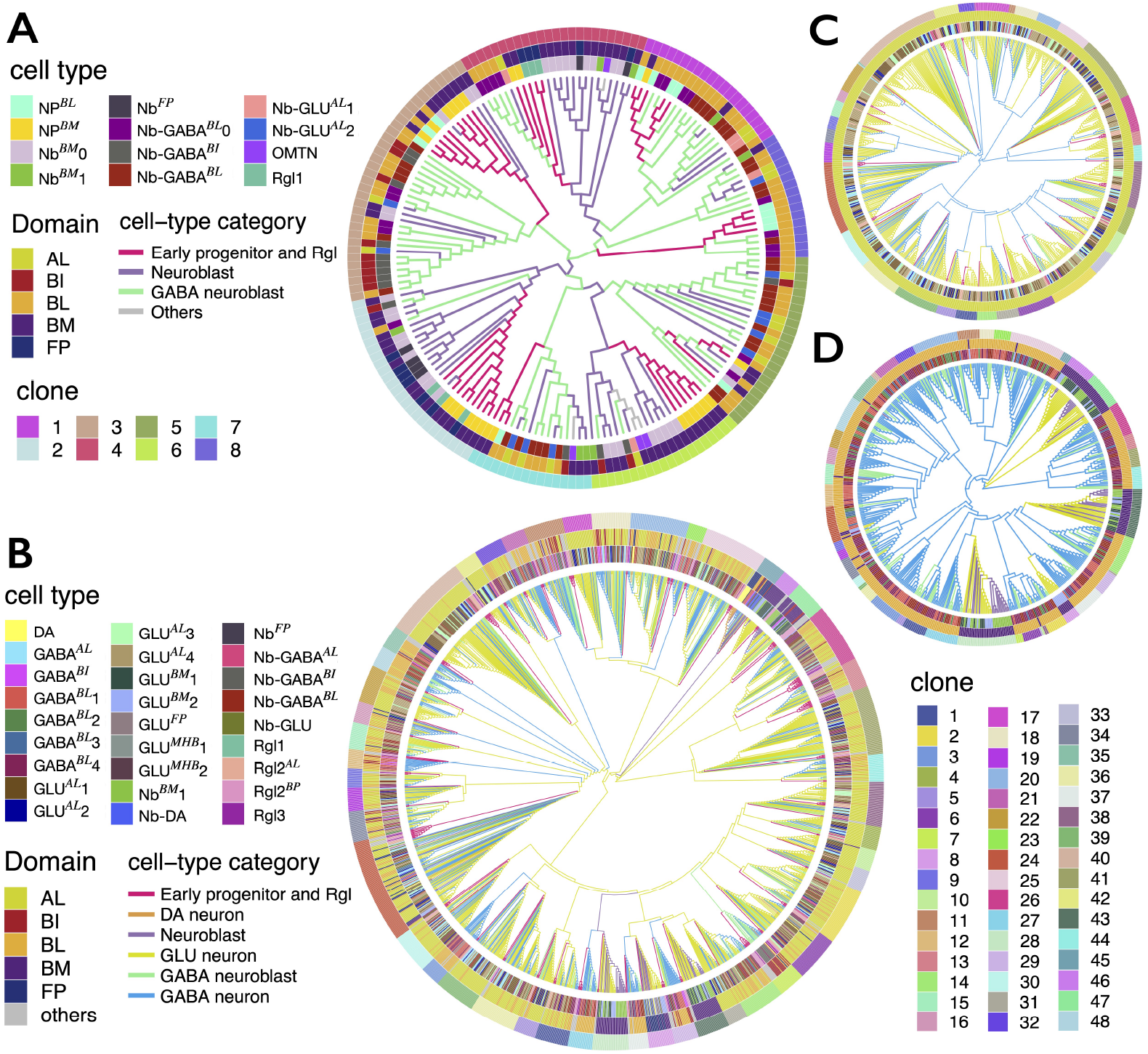
BiLinT analysis of lineage development in the mouse vMB at E11.5 and E15.5. (A) Reconstructed cell lineage tree at E11.5 from one of three biological replicates. The lineage trees for the other two replicates are shown in Fig. S5. (B) Reconstructed cell lineage tree at E15.5. (C, D) Magnified views of subtrees from the lineage tree in (B), highlighting the AL domain in (C) and the BL and BM domains in (D). Cell types observed across E11.5 and E15.5 include: NP (neural progenitor), Rgl (radial glia-like cell), Nb (neuroblast), Nb-GABA (GABAergic neuroblast), Nb-GLU (glutamatergic neuroblast), OMTN (oculomotor and trochlear nucleus), and GABA (GABAergic neuron), dopaminergic neurons (DA), glutamatergic neuron (GLU), dopaminergic neuroblast (Nb-DA). Each phylogenetic tree in (A)–(D) includes three concentric annotation layers—from inner to outer ring—representing cell type, domain, and clone identity. Branches are colored by cell type category.

Beyond transcriptional dynamics, BiLinT reveals spatially structured lineage relationships that reflect region-specific differentiation trajectories within the developing vMB. At E11.5, lineage trees are dominated by early progenitors, Rgl cells, neuroblasts, and GABAergic neuroblasts (Fig. 5A, Fig. S5A–C), while by E15.5, the trees are largely composed of mature GLU and GABAergic neurons (Fig. 5B, Fig. S5D). Within the reconstructed trees, BiLinT identifies subtrees enriched for domain-specific cell types: GLU and GABA neurons in the AL domain (Fig. 5C), GABA neurons in the BL domain, and GLU neurons in the BM domain (Fig. 5D). The FP domain gives rise to both DA and GLU neurons, as shown by clones 8 and 45 (Fig. 5B). These patterns suggest that BiLinT not only captures clonal structure but also resolves how spatial identity and lineage restriction co-emerge during development.

By jointly modeling barcode and gene expression data, BiLinT resolves clonal relationships that are ambiguous under single-modality methods. For example, within the AL domain, BiLinT places GABA^AL^ and GLU^AL^1–4 neurons within the same clone, reflecting a shared clonal origin (Fig. 5C). Barcode-only phylogenies support this relationship, but expression-only trees separate them (Xie et al. 2023). BiLinT’s unified tree reconciles this discrepancy by showing that these subtypes emerge from a common lineage but diverge transcriptionally during differentiation (Fig. 5C). In contrast, GABA^BL^1–4 and GLU^BM^1–2 arise from distinct clones corresponding to BL and BM lineages, respectively (Fig. 5D). BiLinT also identifies a shared lineage between DA neurons and Nb^FP^ in the FP domain, highlighting a common progenitor (Fig. 5B).

### Integration of BiLinT trees reveals clonal dynamics across embryonic time

Having reconstructed lineage trees at each stage with BiLinT, we perform a co-occurrence–based analysis to connect progenitor states at E11.5 with differentiated outcomes at E15.5. To motivate this analysis, we first summarize domain-specific clonal relationships within each stage (Fig. 6B), guided by the single-stage trees in Fig. 5B–D, which highlight both shared and distinct clonal origins across vMB domains. Building on these observations, we then estimate transition flows by computing the probabilities that early progenitor and Rgl cells differentiate into later states based on clone-wise co-occurrence. Specifically, transition probabilities are computed by multiplying the proportion of early progenitors/Rgl cells with the proportion of each terminal state within the same lineage. These probabilities are aggregated across all clones to generate a lineage transition map, which is filtered by a normalized probability threshold of 0.01 (Fig. 6A). We use four neuroblast types (Nb-GABA^BL^, Nb-GABA^BI^, Nb^BM^1, and Nb^FP^), present in both stages, as intermediate states to extend transitions into E15.5. This analysis reveals four major differentiation trajectories: (1) from NP^BL^ to multiple GABA neurons (GABA^BL^1–4) via Nb-GABA^BL^; (2) from NP^BM^ to GABA^BI^ via Nb-GABA^BI^; (3) from NP^BM^ to GLU^BM^1–2 via Nb^BM^1; and (4) from Rgl1 to DA neurons via Nb^FP^ (Fig. 6A,C).

**Figure 6.**
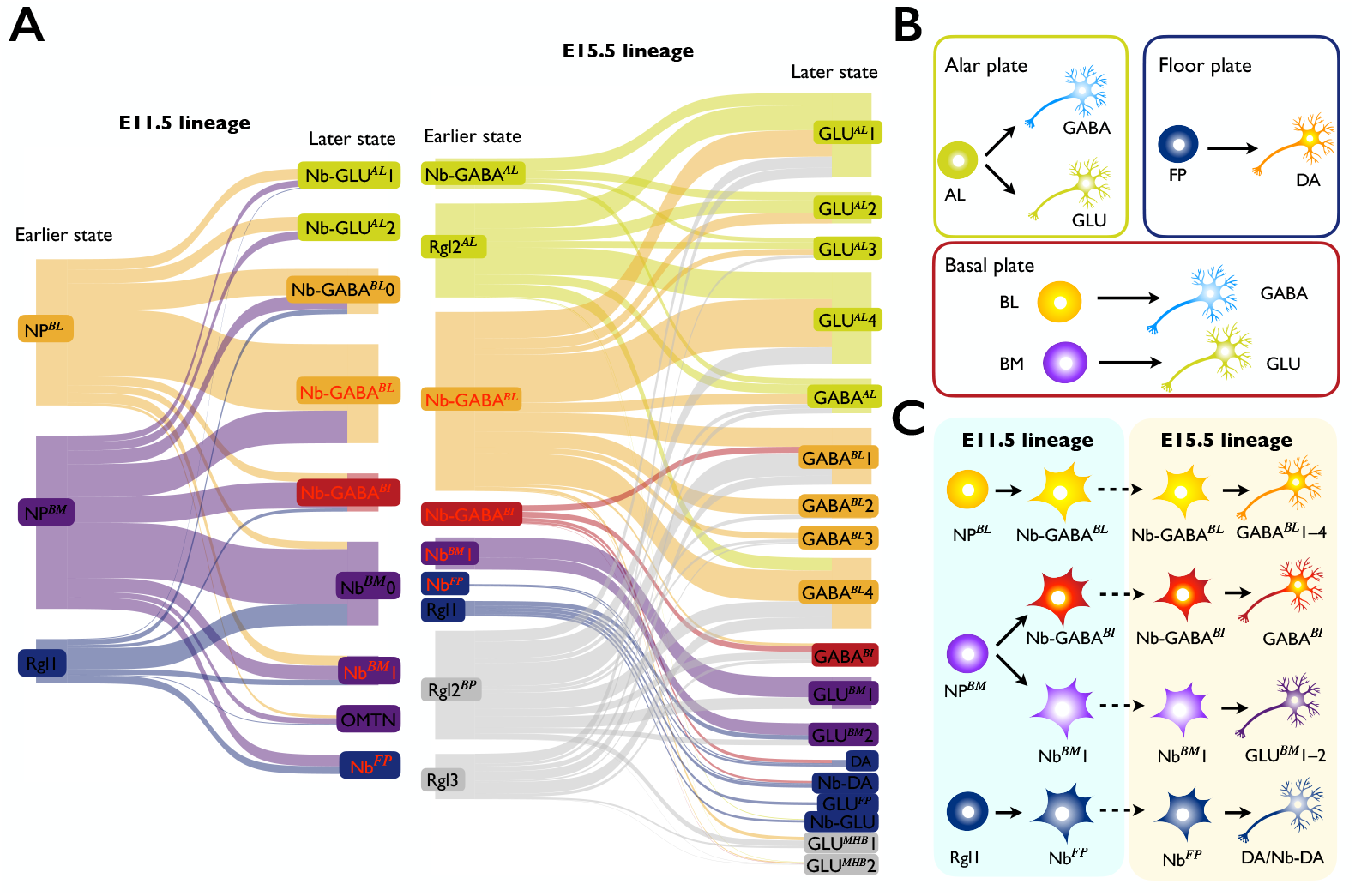
Clone-based transition analysis links progenitor states at E11.5 to differentiated outcomes at E15.5. (A) Sankey diagram showing the transition flow between earlier- and later-state cell types for the E11.5 (left panel) and E15.5 (right panel) time points. (B) Schematic illustration of shared or distinct clonal origins of cell types across different midbrain domains. (C) Major lineage trajectories inferred from (A), tracing the differentiation from early progenitors in the E11.5 vMB through intermediate states to terminal cell fates in the E15.5 vMB.

Together, BiLinT’s within-stage reconstructions (Fig. 5) and this cross-stage co-occurrence analysis resolve clonal origins, developmental trajectories, and domain-specific lineage bifurcations in the developing vMB.

## Discussion

The integration of single-cell multimodal data has revolutionized our understanding of cellular dynamics, particularly in developmental biology and cancer research. The proposed BiLinT framework represents a significant advancement in the reconstruction of lineage trees by incorporating both gene expression and lineage barcode data into a unified probabilistic model. This approach addresses critical limitations of existing methodologies, which often treat lineage and gene expression data separately or rely on fixed phylogenetic structures, thereby neglecting the inherent uncertainties in tree reconstruction.

One of the key strengths of BiLinT is its use of the OU process to model the continuous evolution of gene expression along the branches of a time-scaled Bayesian phylogenetic tree. This allows for a more accurate representation of the dynamic nature of cellular processes, where gene expression levels are influenced by both lineage history and external factors. By integrating gene expression data with lineage barcode information, BiLinT provides a more refined view of cell fate decisions and differentiation pathways, which is critical for understanding complex biological systems.

Our framework assumes conditional independence between barcode-derived lineage information and gene expression profiles, given the phylogenetic tree. This simplifies joint modeling by decoupling lineage dynamics from transcriptomic variation, as CRISPR-induced barcodes are heritable markers rather than direct gene expression regulators. While dependencies may arise from selective pressures or cellular stress, our method captures gene expression evolution along the phylogeny. Future models could address potential feedback between lineage and gene expression. Experimental validation of this assumption will strengthen the framework.

To simplify the model and improve computational efficiency, we assume that the root and stationary covariances are scalar multiples of the identity matrix, and that observational noise is identical across genes. By combining these assumptions with a input of principal component analysis (PCA) top components, we reduce the number of model parameters and computational cost. Importantly, our simulation results suggest that these simplifications introduce negligible systematic bias, while still capturing the overall data structure.

In comparison to existing tools like LinRace and LinTIMaT, which also aim to integrate lineage and gene expression data, BiLinT offers a unified probabilistic approach that explicitly models the uncertainties in tree reconstruction and the continuous nature of gene expression. This represents a shift from traditional two-step phylogenetic inference methods, where the phylogenetic tree is often assumed to be known and fixed before analyzing gene expression data. By contrast, BiLinT simultaneously infers the latent tree structure and the associated parameters, providing a more holistic view of cellular lineage and gene expression evolution.

## Methods

### Model of BiLinT

To jointly reconstruct cell lineage and infer the evolution of cell states, we develop a bi-process modeling framework that describes both barcode mutations and gene expression dynamics along a shared phylogenetic tree 𝒯. These two data modalities evolve independently but coherently under time-scaled stochastic processes. The lineage barcode evolution is modeled via a continuous-time substitution process, while gene expression follows a hierarchical General Gaussian Process, both parameterized with respect to the same tree topology and branch lengths.

We follow the framework of TiDeTree and adopt a Time-scaled Substitution Model to describe barcode evolution on a rooted phylogenetic tree 𝒯 (yellow module in Fig. 1D). The model tracks *N* observed cells, each with *K* genomic targets subject to modifications by editing enzymes. Each target carries a unique barcode, enabling precise differentiation. Initially, all targets are unedited. During a defined scarring window (*t*_1_ *≤ t ≤ t*_2_), targets can transition from the unedited state to one of *S* scarred states due to enzymatic activity. Editing is induced either by injecting enzymes at *t*_1_ > = 0 or by activating enzyme expression later (*t*_1_ *>* 0). The endpoint *t*_2_ is typically inferred from independent experiments identifying when the proportion of unedited targets stabilizes (Alemany et al. 2018; Spanjaard et al. 2018; Seidel and Stadler 2022).

Throughout the experiment, targets may also undergo silencing, a permanent deactivation that prevents further modification. Silencing can occur at any time, whereas scarring is restricted to the scarring window. Once a target is silenced, it can no longer be scarred, and a scarred target cannot transition between scarred states. The final states across all observed cells are recorded in a matrix *Z*, where *Z*_*ik*_ denotes the state of the *k*-th target in the *i*-th cell.

To model these dynamics, TiDeTree employs a continuous-time Markov chain with a state space Ω = {unedited, silenced, scarred}. Each target starts unedited, with transitions governed by a time-dependent rate matrix *Q* as follows.

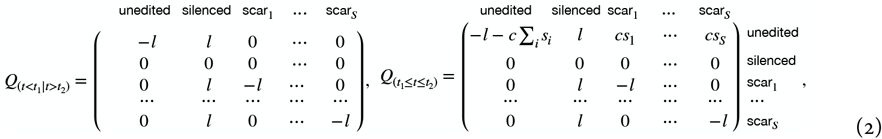

where silencing occurs at a constant rate *l* throughout the experiment, while scarring is restricted to the window [*t*_1_, *t*_2_] with rates ***s*** = {*s*_1_, *s*_2_, …, *s*_*S*_}. Both processes are irreversible: unedited and scarred targets can be silenced at any time; once a target is either silenced or scarred, no subsequent editing occurs.

The model accommodates *S* scarred states, each representing a distinct editing outcome. The editing rate *c* determines the accumulation rate of scarring states over time, with transition rates parameterized as *c* × *s*_*i*_ for each state *i*. Setting *s*_*S*_ = 1 ensures that *s*_*i*_ functions as a rate multiplier, reflecting the relative frequency of each state. Alternatively, *c* can be interpreted as a molecular clock rate, capturing the overall rate of barcode editing events.

While the substitution model is not time-reversible, its transition matrices are diagonalizable, allowing analytical computation of the transition probability matrix *P*_*t*_ (Supplementary Note 1). This enables efficient inference with *O*(*k*) computational complexity, where *k* = *S* + 2 is the dimension of the rate matrix, significantly improving efficiency over the standard *O*(*k*^3^) diagonalization approach. Integrating these principles, TiDeTree enables robust time-scaled inference of lineage barcodes within a phylogenetic framework.

To jointly incorporate transcriptomic information, we employ a General Gaussian Process Model to model how gene expression evolves in the context of a rooted phylogenetic tree 𝒯 with *N* tips and *m* internal nodes (blue module in Fig. 1D). Each observed cell *i* (1 ≤ *i* ≤ *N*) is associated with a vector ***Y***_*i*_ of gene expression in a *p*-dimensional space, while the internal nodes represent latent cell states denoted by ***V***_*j*_ (1 ≤ *j* ≤ *N* + *m*). Each non-root node *j* has a unique parent node *p*(*j*) and an associated branch length *l*_*j*_. The model is built hierarchically to describe both how the root state initializes the system and how cell states propagate along the branches of the tree.

To capture the dynamics of cell state evolution, we assume that transitions along the tree follow a continuous-time OU process (Bastide et al. 2021). The OU process introduces drift toward an optimal value over time, ensuring that variance remains bounded—unlike Brownian motion, which accumulates variance indefinitely. This approach allows us to reflect the stochastic nature of gene expression while still accounting for the selection strength that drives convergence toward stable expression levels. Below, we outline the components of the General Gaussian Process Model used to describe these transitions.

Root Model. The state at the root of the tree, ***V***_*r*_, initializes the propagation process. It is drawn from a multivariate normal distribution with mean ***µ*** and variance **Γ**:

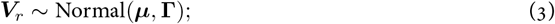

This step defines the initial state of the system, reflecting the expected mean expression values and their variability at the root of the phylogenetic tree. For simplicity, we assume **Γ** = *σ*_*r*_***I***.

Propagation Model. For each non-root node *j*, the latent state ***V***_*j*_ depends on the state of its parent node ***V***_*p*(*j*)_. The state transition follows a Gaussian distribution with mean determined by the parent’s state and parameters reflecting drift and variability along the branch:

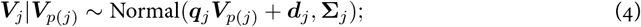

Here, the terms are defined as: (1)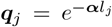, controlling how much the state decays over the branch length *l*_*j*_, where ***α*** governs the selection strength of the OU process; (2)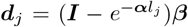, representing drift toward the optimal value ***β***; (3)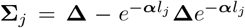, representing the variance structure over the branch, where **Δ** is the stationary variance of the OU process. To reduce noise and dimensionality, we apply PCA to the original gene expression matrix and use the top principal components as the input features for modeling. This choice transforms the gene expression data into a lower-dimensional space that captures the dominant modes of variation while reducing the effects of technical noise. In this PCA basis, the coordinates are orthogonal and uncorrelated, making a diagonal stationary covariance for the OU process natural. Therefore, we assume **Δ** = *σ*_*p*_***I*** for simplicity, where *σ*_*p*_ controls the extent of process variance in the PCA space. This model captures the balance between random fluctuations and drift toward a stable expression state, reflecting how expression levels evolve across cell generations.

Observation Model. Each observed gene expression vector ***Y***_*i*_ corresponds to the latent state of the parent node ***V***_*p*(*i*)_ with an additional layer of observational noise:

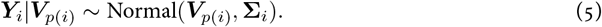

We assume the observational noise to be identical across genes, simplifying the covariance matrix to **Σ**_***i***_ = *σ*_*o*_***I***. The hierarchical structure can be reformulated as a unified view of the propagation process (Supplementary Note 2), treating both latent states and observations in a consistent manner.

The model allows for multiple measurements ***Y***_*i*_ to be associated with the same latent state ***V***_*p*(*i*)_, enabling the construction of a clone tree (Fig. S6). In the clone tree, multiple observations arise from a single stable state, each with variance *l*_*i*_*σ*_*o*_, where *l*_*i*_ reflects the cell-type-specific variability. This contrasts with the simpler variance structure of the cell tree, where all observations share the same noise variance *σ*_*o*_. The refined subpopulations of stable cell types in the clone tree are derived by clustering cells sharing the same barcode state (see Supplementary Note 3). Accounting for potential measurement variability across multiple instances of the same cell type, the clone tree framework provides a realistic modeling of biological systems.

This Gaussian process framework, illustrated by the blue module in Fig. 1D, captures the evolution of gene expression along the phylogenetic tree. By leveraging the OU process and incorporating observational noise, it provides a flexible yet structured approach to modeling complex cell state dynamics.

### Inference of BiLinT

We employ the MCMC method to estimate the unknown parameters in BiLinT, primarily utilizing the Metropolis–Hastings (MH) algorithm for parameter sampling. We detail our sampling procedures, covering the likelihood computation, prior distribution assignment, initialization, and the proposal of new values within the MH sampling framework for all parameters for both cell and clone tree in Supplementary Note 3).

To prevent the Markov chain from becoming trapped in local optima, we employ the parallel tempering technique. This approach involves running multiple chains at different temperatures and allowing exchanges of samples between them (Z Chen et al. 2022). For each parallel chain, we initialize the phylogenetic tree 𝒯 using the heuristic method described in Supplementary Note 3 and set the remaining parameters according to their respective prior distributions before commencing MCMC sampling. We then sample from the posterior *p*(*ϕ*, 𝒯, ***φ***|***Y***, ***Z***) via MH moves and report the maximum a posteriori (MAP) estimate, defined as the sampled state with the largest posterior value.

### Benchmarking BiLinT

#### Simulation data

To systematically evaluate BiLinT’s ability to integrate single-cell RNA-seq and lineage barcode data, we design three simulation settings and compare BiLinT with other state-of-the-art methods.

Simulation 1 is based on the TedSim simulator (Pan, H Li, and X Zhang 2022; Pan, H Li, Putta, et al. 2023), which uses an asymmetric cell division model to simulate gene expression along a balanced lineage tree. Barcode data are generated using our continuous-time, time-scaled substitution model. We vary one parameter at a time—clock rate (*c*), silencing rate (*l*), or number of cells (*N*)—while keeping the others fixed. Each parameter setting includes 10 replicates, resulting in 90 paired datasets. Default values are *N* = 64, *c* = 5 × 10^−4^, and *l* = 0.01.

Simulation 2 extends the evaluation to unbalanced trees sampled from a birth–death–sampling process. We use our General Gaussian Process Model to simulate gene expression and the same substitution model for barcode data. This scenario includes 30 datasets, varying only the number of cells, with model parameters drawn from the priors described in Supplementary Note 3.

Simulation 3 explores a challenging case in which barcode data alone are insufficient to resolve lineage structure due to limited target sites and high parallel mutation rates. Eight clones are simulated, each with eight cells, arranged along an unbalanced tree (Fig. 3A). Barcode data (Table S1) provide weak phylogenetic signal, whereas gene expression patterns are strongly informative. This design allows us to evaluate BiLinT’s ability to leverage multimodal data for accurate reconstruction.

#### Real data

In addition to simulation data, we also apply BiLinT to five real single-cell lineage-tracing datasets. The first four datasets consist of temporal single-cell lineage-tracing data from the developing mouse vMB, including three biological replicates at E11.5 and one at E15.5, obtained using the snapshotting CREST (snapCREST) method (Xie et al. 2023). These mouse developing vMB data are available under NCBI GEO accession GSE210139. The fifth dataset is generated from developing HSCs using DARLIN, an inducible barcoding system that enables lineage tracing and analysis across mouse tissues, as well as combined transcriptional single-cell measurements (L Li et al. 2023). The mouse hematopoietic development data are available under GSE222486.

#### Evaluations

We compare the performance of our algorithm against four recent methods for analyzing single-cell lineage-tracing data: LinRace (Pan, H Li, Putta, et al. 2023), LinTIMaT (Zafar et al. 2020), Cassiopeia (Jones, Khodaverdian, et al. 2020), and DCLEAR (Gong, Kim, et al. 2022). Our evaluation employs three metrics: Robinson-Foulds (RF) distance (Robinson and Foulds 1981; Steel and Penny 1993), Subtree Prune and Regraft (SPR) distance (Oliveira Martins et al. 2008; De Oliveira Martins et al. 2016), and Path distance (Steel and Penny 1993). Details of the distance metrics are in Supplementary Note 4.

## Software availability

The BiLinT software is publicly available at https://github.com/ucasdp/BiLinT.

## Data access

All simulation data generated and analyzed during the current study are available from the corresponding author on reasonable request.

## Acknowledgements

We thank the Yuejun Chen lab (Institute of Neuroscience, CAS, Shanghai) for providing preprocessed code and data for the mouse vMB analyses used in this work. This work was supported by National Key Research and Development Program of China [grant number 2022YFA1004801].

Authors’ contributions: ZC, LW, and LM conceived the model. ZC and BZ developed the code and conducted the experiments. ZC, BZ, LT, LW, and LM analyzed the results. ZC, BZ, LT, FG, LW, and LM wrote and reviewed the manuscript. All authors read and approved the final manuscript.

